# TreeTerminus - Creating transcript trees using inferential replicate counts

**DOI:** 10.1101/2022.11.01.514769

**Authors:** Noor Pratap Singh, Michael I. Love, Rob Patro

## Abstract

The accuracy and robustness of many types of analyses performed using RNA-seq data are directly impacted by the quality of the transcript and gene abundance estimates inferred from this data. However, a certain degree of uncertainty is always associated with the transcript abundance estimates. This uncertainty may make many downstream analyses, such as differential testing, difficult for certain transcripts. Conversely, gene-level analysis, though less ambiguous, is often too coarse-grained. To circumvent this problem, methods have proposed grouping transcripts together into distinct inferential units that should be used as a base unit for analysis. However, these methods don’t take downstream analysis into account.

We introduce `TreeTerminus`, a data-driven approach for grouping transcripts into a tree structure where leaves represent individual transcripts and internal nodes represent an aggregation of a transcript set. `TreeTerminus` constructs trees such that, on average, the inferential uncertainty decreases as we ascend the tree topology. The tree provides the flexibility to analyze data at nodes that are at different levels of resolution in the tree and can be tuned depending on the analysis of interest. To obtain fixed groups for the downstream analysis, we provide a dynamic programming (DP) approach that can be used to find a cut through the tree that optimizes one of several different objectives.

We evaluated `TreeTerminus` on two simulated and two experimental datasets, and observed an improved performance compared to transcripts (leaves) and other methods under several different metrics.

## 1 Introduction

Transcript abundance estimation is among the key target applications of RNA-Seq. The variation in transcript expression plays a key role in development [3, 40, 28]; and characterization of diseases and their subtypes [41, 46, 35]. One of the approaches towards estimation of transcript abundances is to probabilistically assign a given fragment to the transcripts using maximum likelihood or Bayesian inference [24, 6, 30]. This has to be done since there exists an ambiguity towards finding the true locus of origin for a given sequencing fragment, when it can map equally well to the shared sequences within transcripts. Thus, a certain degree of uncertainty is associated with the transcript abundance estimates, depending on the nature of the fragments. This in turn makes downstream analysis, such as differential testing, difficult for certain transcripts. Uncertainty also exists for gene expression estimates but is less compared to transcripts, since there will be less ambiguity when mapping fragments to genes. The uncertainty of a transcript/gene can be estimated from the inferential replicates generated either through MCMC/Gibbs Sampling; or through bootstrap sampling of the reads [24, 30, 44, 15, 6]. The highly uncertain transcripts/genes may become invisible when the inferential replicates are incorporated into the downstream tasks To circumvent this problem, some methods have grouped transcripts/genes into distinct inferential units that share a lot of multi-mapping reads [32]. `mmcollpase` [43] proposes to group transcripts by computing correlation between every pair of transcripts on the posterior replicates and the pair that has the most negative correlation is grouped. This process is repeated for multiple iterations until a stopping criterion is reached, where at each iteration the correlation is recomputed for every transcript/group with the group from the previous iteration and a new group is formed. `Terminus` [33] also provides transcript groups as an output but employs difference in inferential relative variance [48] between the group and the mean of its underlying transcripts/subgroups as the statistic for group creation. It first creates a graph on the transcripts, with an edge denoting that the transcripts co-occur in at least one `range-factorized` equivalence class [47] and only the transcripts that are connected by a path on the graph are considered for grouping. `Terminus` first finds groups across each individual sample and uses a consensus approach to output consensus groups across samples. Importantly, both `mmcollapse` and `Terminus` provide a single level of resolution for analysis, not permitting analysis across different levels.

Another limitation with both of the above methods, is that the resolution at which grouping is stopped is governed by a threshold determined heuristically that does not take the downstream analysis into account. When performing downstream tasks such as differential analysis, these methods can either thus over-aggregate transcripts, masking the signal in the process or under-aggregate leading to weaker signals, and an inability to make confident calls. Since the goal of these methods is to provide concrete groups as an output without doing much aggregation, they introduce many filtering constraints on transcripts in order for them to be considered for aggregation which might further lead to missing good candidates for aggregation.

In this work we introduce `TreeTerminus`, a new method that aims to address some of the above mentioned shortcomings. `TreeTerminus` expands upon the idea of `Terminus` and groups transcripts in a tree structure where leaves represent individual transcripts and internal nodes represent an aggregated set of leaf transcripts. Across the samples in the experiment, inferential uncertainty decreases on an average, ascending the tree topology. `TreeTerminus` can be run on a single sample and we provide two different approaches in order to extend it for the multi-sample settings. The tree provides the flexibility to analyze data at nodes that are at different levels of resolution in the tree and can be tuned depending on the analysis of interest. To obtain fixed groups for downstream analysis, we provide a dynamic programming (DP) approach that can be used to find a cut through the tree that optimizes one of several different objectives. In addition, the DP approach has the ability to optimize for other user-defined objective functions provided they adhere to the required constraints necessary to be efficiently optimized on a tree.

To the best of our knowledge this is the first time that transcripts have been arranged in a tree-like structure, that too in a data-driven manner. We evaluated `TreeTerminus` on two simulated and two experimental datasets, and observed an improved performance compared to transcripts (leaves) and other methods under several different metrics. `TreeTerminus` has been implemented in Rust and the source code can be downloaded from https://github.com/COMBINE-lab/TreeTerminus. We have also created an R Package `beaveR` that parses the output of `TreeTerminus`, implements DP algorithms for finding an optimal cut, and provides helper functions to obtain useful statistics for subtrees within the `TreeTerminus` -derived tree structures. The `R` Package can be downloaded from https://github.com/NPSDC/beaveR.

## 2 Methods

We first briefly describe `Terminus` [33], as our method builds on top of it. The `group` step of `Terminus` takes as input an individual `Salmon` [30] quantified sample from an RNA-Seq experiment and outputs a set of transcript groups. It makes use of the `range-factorized` equivalence classes *ξ* [47], and inferential replicates 𝒫 obtained after running `Salmon` with the appropriate flags. An equivalence class denotes an association from a set of transcripts to a set of reads, that are mapped to all the transcripts in that set. A `range-factorized` equivalence class in addition also encodes the mapping quality, with a single class constituting a set of pairs (*t*_*i*_,*w*_*i*_) rather than just set of *t*_*i*_, where *t*_*i*_ denotes the transcript and *w*_*i*_ represents the average conditional probability with which the fragments in the equivalence class arose from that transcript. The `Terminus` groups consist of transcripts that share large numbers of ambiguously-mapped fragments. It employs a union-find data structure, with the first step being to scan over *ξ* and group the transcripts that appear in the same set of equivalence classes and have near-identical conditional probability vectors using the `Union` operation. This ensures that these transcripts will belong to the same partition, when checked through the `Find` operation. It then constructs a graph with the transcripts as nodes, where an edge between any two nodes *v*_*i*_,*v*_*j*_ implies that they co-occur in at least one equivalence class and have the edge score *s*(*v*_*i*_,*v*_*j*_) ≤*τ* where

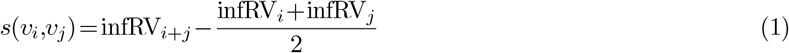

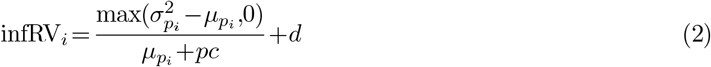

*p*_*i*_ are the posterior (`Gibbs`) replicates for transcript *i*, 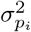 and 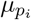 are the variance and mean over the posterior replicates, *pc* is the pseudocount (default is 5), *d* is a small global shift (default is 0.01) and *τ* is a threshold of difference in inferential relative variance. A min heap `H` is constructed over the edges keyed by *s*(*v*_*i*_,*v*_*j*_) and at each iteration *t* an edge with the lowest score is popped out till the iteration `H` becomes empty. If the partition(group) of any endpoint vertex for the popped edge has been modified but its corresponding score was not updated, then the edge is called stale. If the edge is not stale then the corresponding nodes *v*_*i*_,*v*_*j*_ are grouped using the `Union` operation and the posterior samples for *v*_*i*_ are updated with *p*_*i*_ = *p*_*i*_ +*p*_*j*_, assuming *i<j*. The score *s*(*v*_*i*_,*x*) is recomputed for all the vertices {*x*|*x* ∈ *adj*(*v*_*i*_) ∪ *adj*(*v*_*j*_) − {*v*_*i*_,*v*_*j*_}} with *adj*(*v*_*i*_) denoting the neighbours of vertex *i* in the graph at iteration *t*. The edge with the updated *s*(*v*_*i*_,*x*) is pushed to `H`, if *s*(*v*_*i*_,*x*) ≤ *τ*. The `consensus` step outputs a set of transcript groups across samples using the groups obtained for the individual samples (see Section S1.1).

### 2.1 TreeTerminus

The transcripts in a `Terminus` group consist of at least two transcripts that have reads multi-mapped to them, which is the source of uncertainty for transcript abundance estimation. `TreeTerminus` outputs a tree for each individual group, encoding the summarized order in which a set of transcripts should be aggregated within a group, such that across the samples, average uncertainty decreases as we ascend the tree created for that group.

`TreeTerminus` has a `group` step, that constructs transcript-trees for a single sample. It starts by creating a single node tree corresponding to each transcript. For any pair of transcripts/subgroups that are aggregated in the `group` step of `Terminus`, a binary tree is created with its children being the individual trees corresponding to the node-pairs that are grouped. The process of tree creation continues till the time heap `H` becomes empty. In addition, some of the constraints that `Terminus` imposed on transcripts/groups before they could be even considered for further grouping have also been relaxed. It is no longer necessary that a node *v*_*i*_ should have *infRV* (*v*_*i*_)≥*δ* (called filtered or `Filt`) and that for a pair of nodes *v*_*i*_,*v*_*j*_, *s*(*v*_*i*_,*v*_*j*_) ≤ *τ* (called early stop or `ES`). For the *N* groups that will be obtained from a sample *m*, a set of *N* trees *T*^*m*^ = {*T*_1*m*_,…,*T*_*Nm*_} will be generated, such that for a group *g, Λ*(*g*) = *Λ*(*T*_*gm*_), with *Λ* representing the set of transcripts covered by a tree or a group and *Δ*^*m*^ represents all the trees belonging to sample *m*. We next propose and describe two different approaches to obtain trees, representative of all samples in the experiment.

#### Mean Tree

This is a single step procedure where all the samples are processed together and a single tree is produced from all the samples w.r.t a group. The steps in the tree construction follow directly from what we have described above, with some modifications. To find transcripts that have near identical conditional probability weights across all the equivalence classes, the search space is expanded to all samples rather than a single sample. Later, when the graph across transcripts is constructed, an edge between any two nodes implies that they co-occur in the same equivalence class in at least one sample where the edge score *s*^′^(*v*, *v*) is updated as, 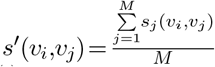, and *M* represents the total number of samples in the experiment. The indicator function *h*(.) that determines whether a transcript is a good candidate for aggregation when considered in isolation is also updated as:

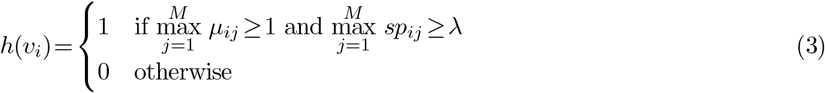

where *μ*_*ij*_ =mean(*p*_*ij*_) and 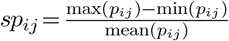, with *p*_*ij*_ denoting the posterior replicates for transcript *i* in sample *j* and *λ* is a user-defined parameter that is set to 0.1 that serves as a threshold for spread.

### 2.2 Consensus tree

Construction of consensus trees is more involved and takes as input the sample group trees obtained by running `group` step on each sample. Given a set of trees for a given transcript group across samples, we want to find a tree that summarizes the topological structure of the input trees. Such a tree is called consensus tree [1, 19]. There exists a wide variety of methods to get consensus trees, depending on the input and downstream applications [8, 27, 19, 7]. In this work we have used the majority rule extended or greedy consensus tree algorithm[8, 11], and is described in Section S1.2.

A constraint of a consensus tree algorithm is that it requires all the input trees should span the same leaf set. To obtain a consensus tree *T*_*g*_ for group *g* in an experiment containing *M* samples, we want to provide group trees across samples *Δ*_*g*_ = {*T*_*g*1_,…,*T*_*gM*_} as an input to the Majority Rule Extended Consensus Tree Algorithm, with *T*_*gi*_ representing the tree for sample *i* for the group *g* and *Λ*(*g*) = *Λ*(*T*_*gi*_), ∀ *i* ∈{1,…,*M*}. *Λ* represents the transcripts covered by a tree or a group. However, a group covering a transcript set in one sample might not be preserved in other samples. We can have tree/s within a sample that covers a transcript set that overlaps but is not the same as the transcript(leaf) set in a tree from some other sample in the experiment. This is demonstrated in the example shown in Figure 1a. For the five different samples, we have trees that cover different transcript sets, covering overlapping transcripts. First sample contains the tree built on 5 transcripts. The second and fifth samples span the same set of 4 transcripts but are subsets of the transcripts covered by *T*_11_. The third and fourth samples each have two trees containing transcripts that are also a subset of *Λ*(*T*_11_). Thus, we cannot directly apply the consensus tree algorithm on the trees obtained by the `group` step of `TreeTerminus`.

**Fig. 1.**
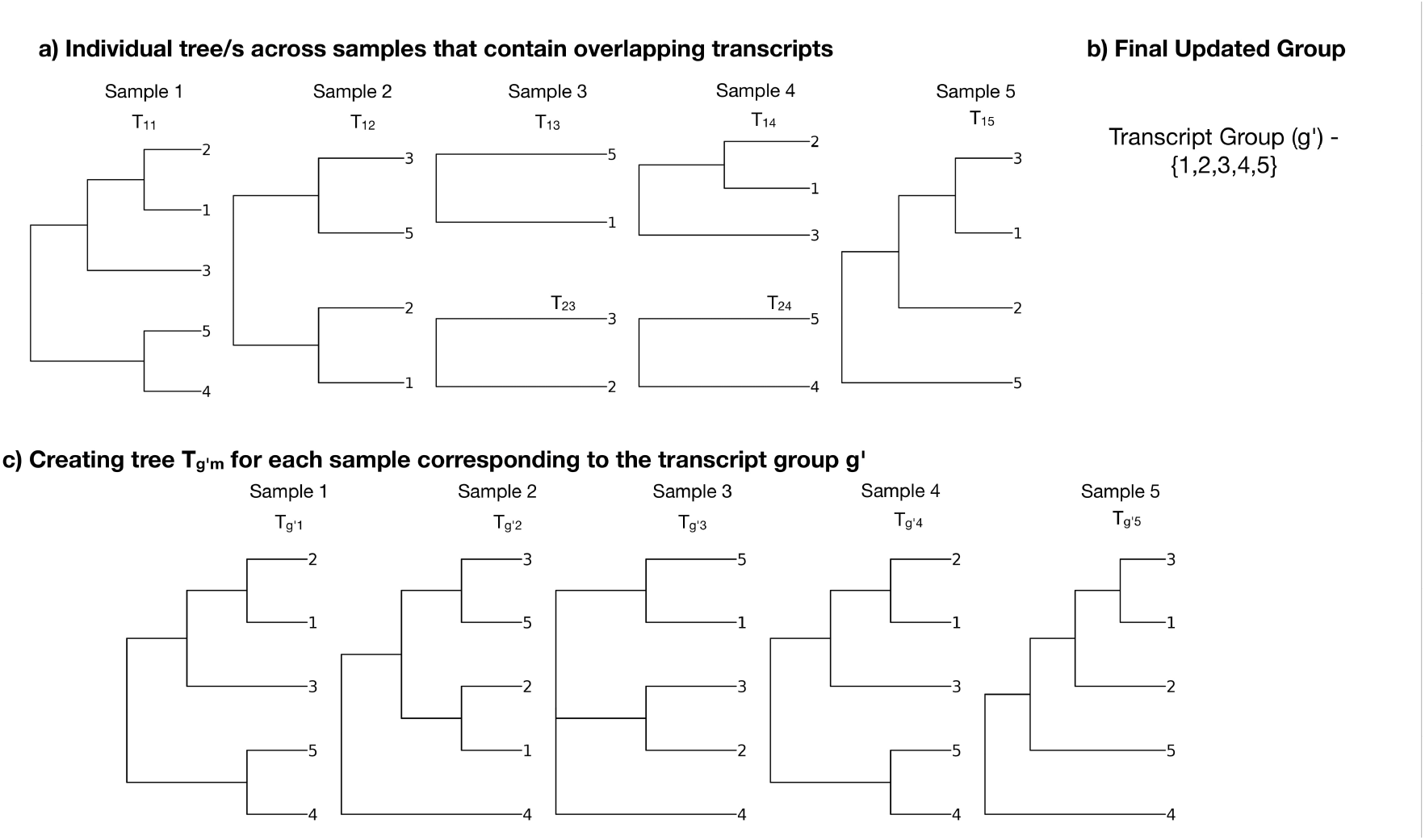
Example of a sample group to demonstrate why consensus step cannot be directly applied on the trees obtained from `group` step for samples and the modifications made in order to apply the consensus tree algorithm. **a** Trees across the 5 different samples that contain overlapping transcripts. Each tree is labelled as *T*_*gm*_, where *m* denotes the sample and *g* denotes a group in that sample. **b** Updated group that is a superset of all the transcripts in the individual trees across samples. **c** Tree created for each sample w.r.t updated group *g*^′^, with 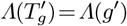.

To resolve this issue, we thus first create a set of updated groups 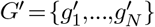, with updated group 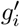 representing a union of all transcripts for all trees across samples that contain overlapping transcripts. Further, the transcripts covered by any such tree should not contain any overlapping transcripts with the transcripts of any other updated group 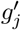 or:

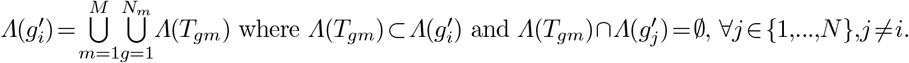

To create the updated group 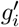, we employ the union-find data structure. We scan through the trees across all samples and apply `Union` operation on the transcript set covered by the tree, which will group these transcripts covered. This ensures that the transcripts belonging to any two trees *T*_*i*_,*T*_*j*_ across the samples where *Λ*(*T*_*i*_)∩*Λ*(*T*_*j*_)≠ ∅ will be grouped together. For the toy example in Figure 1b, this leads to creation of the updated group spanning transcript set {1,2,3,4,5}. The consensus tree 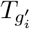 will be created for the updated group 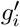, with 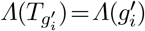.

We next create trees 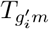 for every sample *m* ∈{1,…*M*} w.r.t each updated group 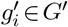, such that tree 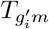 covers same set of transcripts as the updated group or 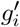 or 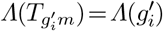. Tree 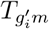 on the updated group 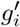 for sample *m* is created using the following steps:

1. For each sample *m*, all the trees *T*_*gm*_ from *Δ*^*m*^ which cover transcripts that overlap with the transcripts covered by 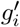 are extracted to form the set 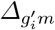 where 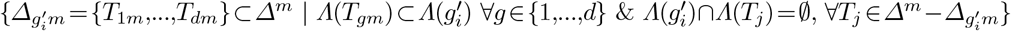.
2. A transcript set `D` is constructed that consists of transcripts covered by the updated group 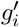 but not by the trees in 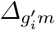. Formally 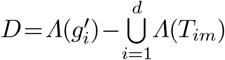, with 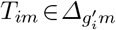.
3. If the set 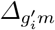 consists of only one tree *T*_*gm*_ and `D` is empty, it implies that the tree *T*_*gm*_ already covers the same transcript set as 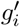 and thus we can safely output *T*_*gm*_ as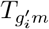.
4. Otherwise, an empty tree is constructed and added to it’s children are all the trees in 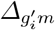 along with the transcripts in the set `D`. This newly created tree forms 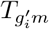.

For each group 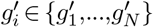, there now exists a set of *M* trees 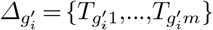, - one for every sample. PHYLIP’s [13, 12] implementation of Majority Rule Extended Consensus Algorithm is then applied on the set 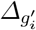 to get the consensus tree 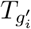 for the updated group 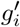. The motivation behind using the majority rule extended consensus algorithm is to provide a chance to preserve a topology present in set of samples that represent a phenotype not present in the majority of samples, which would have been ignored otherwise.

### 2.3 Creating a unified tree from the output of TreeTerminus

Given the output of `TreeTerminus` 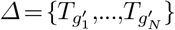, a unified tree *T*_*U*_ is created such that its children consist of all the trees in *Δ*. This is done through our `R` package `beaveR`, which also provides the option to append transcripts to the children of *T*_*U*_ that are not covered by the trees in *Δ* but still should be considered for downstream analysis.

### 2.4 Solving objective functions to obtain discrete inferential units

In order to do any downstream analysis and interpret its results, we require discrete inferential units. These can be obtained by optimizing for an objective. We provide a dynamic programming (DP) approach that can used to optimize different objective functions on a tree, following certain constraints. The DP solving a given objective function outputs a set of nodes in the tree *T* or cut *C* =(*c*_1_,…,*c*_*d*_), where *C* has the following properties:

– The union of leaf nodes belonging to nodes in the cut should cover all the leaf nodes in the tree *T* or 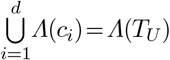.
– The intersection of leaf nodes belonging to any two distinct nodes in the cuts should lead to an empty set or *Λ*(*c*_*i*_)∩*Λ*(*c*_*j*_) =∅,∀*i,j* ∈{1…*d*}.

The DP optimizes the objective functions of the following form to obtain the optimal value and a cut *C*:

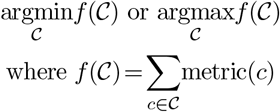

where *metric*(*c*) represents the metric for node *c* in the tree *T*, whose sum we want to optimize over all nodes in the cut. We solve for *f*(*C*) using the procedure described in Algorithm 1. For a given metric of interest, it outputs the optimal value for the objective function and a set of nodes (cut), on which summing up the metric provides the optimal value. Algorithm 2 returns the optimal value for the objective function at a given node and Algorithm 3 returns the cut for a tree/subtree for that obejctive function.

We have solved for the objective functions employing two different metrics to obtain cuts on the tree *T*_*U*_ which we describe in the section below.

#### Algorithm 1

Finding optimal value of the objective function and the corresponding cut for a given metric

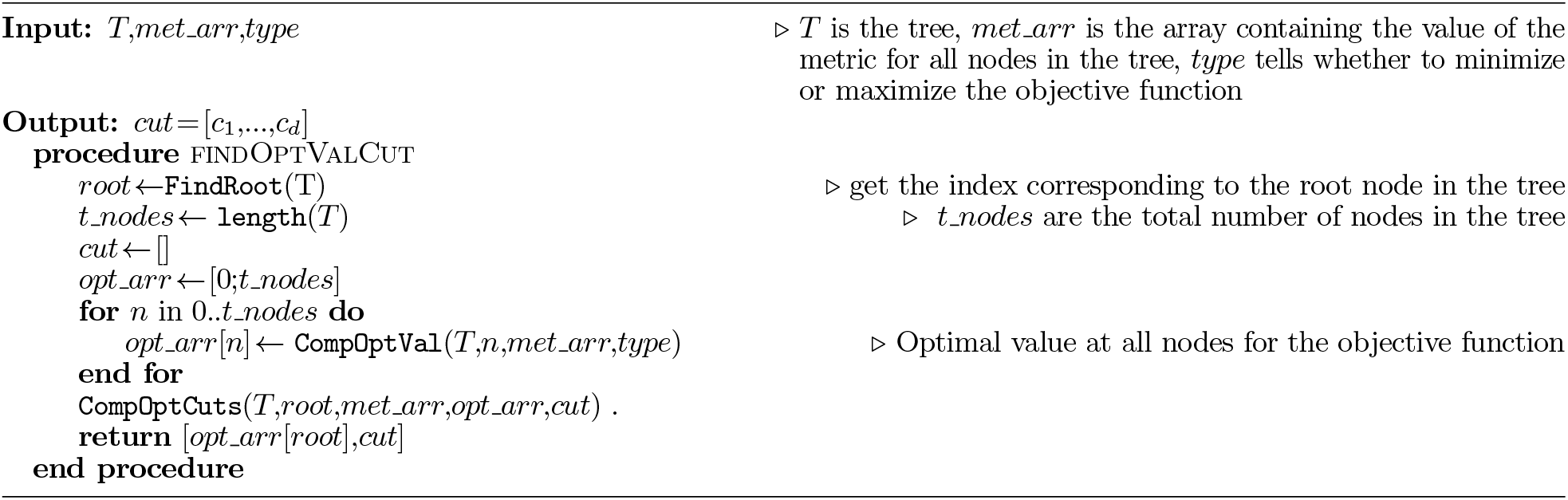

#### Algorithm 2

Function for finding optimal value of objective function at given node

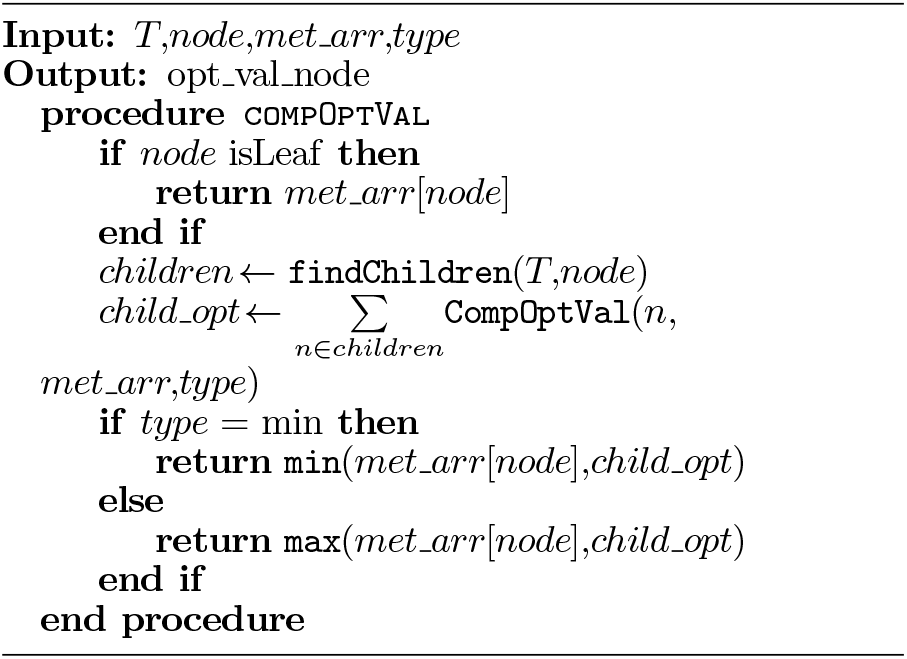

#### Algorithm 3

Function for finding optimal cuts

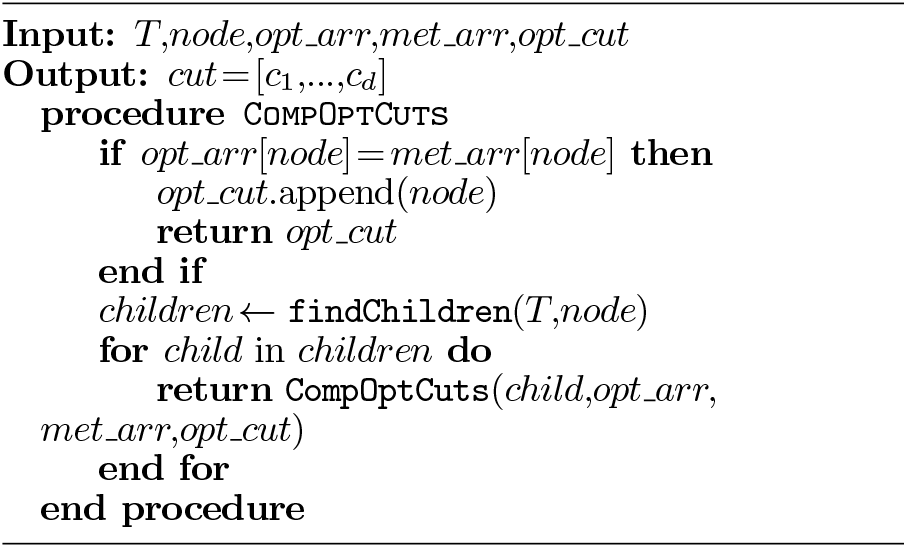

#### Minimize mean infRV and height

For the first objective function, we want to find nodes that have a low mean inferential relative variance and at the same time are at a level close to the leaf in order to provide the finest resolution for downstream analysis. Therefore, in the objective function we minimize the sum for the node metric - sum of mean inferential relative variance and height weighted by *γ*; and then multiplied by the number of descendant leaf nodes. This metric is called `irv_height_desc` and is described formally below with the objective function as:

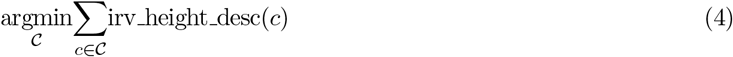

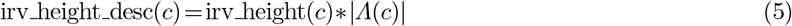

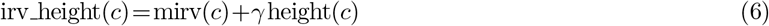

mirv(c) is the mean inferential relative variance for a node across samples, height(*c*) is defined as the number of edges on the longest path from node *c* to its descendant leaf, *Λ*(*c*) represents the descendant leafs for a given node *c*, with *Λ*(*c*) =1, if *c* is a leaf node. The reason that the number of descendants of a node are multiplied is because the value of the objective function is dependent on the number of nodes in the cut and will be biased towards nodes with higher height as they end up replacing multiple lower height descendant nodes. To get the optimal value and desired cut for the above objective function we use the Procedure defined in Algorithm 1 as:

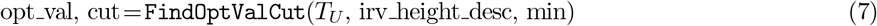

#### Maximize Weighted log fold change

For the second objective function, we want to find nodes that have a high log fold change but a low mean inferential variance. In this objective function, we thus maximize the sum of node metric - absolute log fold change weighted by mean inferential variance multiplied by the size of the descendant leaves for that node. This metric is called `wlfc_desc` and is described formally below with the objective function as:

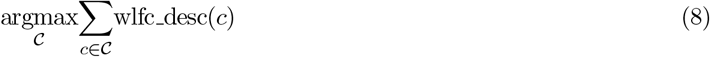

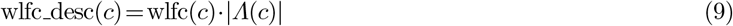

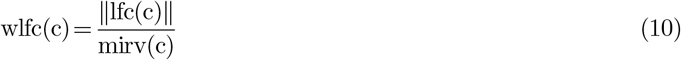

where `lfc` is the log_2_ fold change. Assuming the samples can be grouped into two conditions represented by the sets `m` _1_ and `m` _2_, `lfc` is given by:

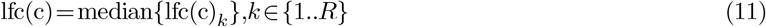

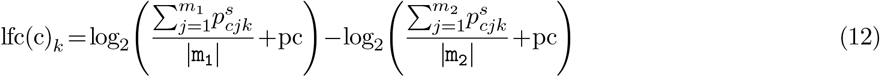

where *R* represents the total number of inferential replicates, 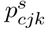 represents the scaled count for sample *j*’s inferential replicate *k* for node *c*, obtained by dividing the count of inferential replicate’s transcript by the depth of inferential replicate. We again use the procedure defined in Algorithm 1 to get the optimal value and the cut as:

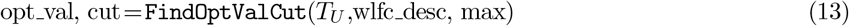

## 3. Experimental Setup

### 3.1 Creation of a baseline anti-correlation tree

For comparison with `TreeTerminus`, we are unaware of any other existing methods that creates transcript-trees. Thus, to create a baseline tree, we create a tree using anti-correlation (negative correlation) between transcripts computed on the inferential replicates, which we also refer to as the `AC` tree. The motivation for using anti-correlation comes from `mmcollapse` [43] which was also used as a baseline method for comparison in `Terminus` [33]. Since `mmcollapse` does not create trees, we use `UPGMA` [37]. `UPGMA` applies a hierarchical clustering procedure on the leaf set to generate a tree, which based on the current input, will group anti-correlated set of transcripts, leading to a reduced uncertainty. We thus believe UPGMA tree is a fair baseline for comparison with the `TreeTerminus` trees, where uncertainty decreases as we ascend the tree.The exact details of constructing the `AC` tree are provided in Section S1.3.

### 3.2 Datasets and quantification pipeline

We ran `TreeTerminus` on two simulated and two experimental bulk RNA-Seq datasets [42, 39]. TPM estimates extracted from GTEx V8 frontal cortex dataset were used as an input to generate the simulated datasets, containing 12 samples in 2 conditions with 6 samples in each. Two variations of the simulated data were created and are referred to as `BrSimNorm` and as `BrSimLow` respectively. The first experimental dataset is derived from two different mouse muscle tissues, consisting of 6 samples in each and we refer it to as `MouseMuscle`. The second experimental dataset is obtained for the brain tissues of Chimpanzee, consisting a total of 73 samples. The two groups for our analysis consist of 5 and 68 tissues respectively. The datasets are described in detail in Section S1.4. `Salmon` was used to quantify the reads from the different datasets, and `Gibbs` sampling was used to generate the inferential replicates. The quantification pipeline is described in Section S1.5.

### 3.3 Different tree methods that have been compared in this study

In this study, we have used both the variations of `TreeTerminus`, with the unified tree obtained after running the procedure in Section 2.3 on the consensus and mean trees referred to as `Cons` and `Mean`. We evaluate their performance on several parameters. To see if there are any benefits of removing the constraints imposed in `Terminus`, we also compare with the unified trees obtained after running the consensus algorithm on the sample trees that were constructed with the variations of those constraints. In the first variation, the consensus tree approach is applied on trees obtained using both constraints i.e `Filt` and `ES` which is referred by `ConsFiltES`. In the second variation, we only apply the constraint `Filt` and the corresponding consensus tree that is created is referred by `ConsFilt`. The Anti-Correlation (`AC`) tree serves as the baseline tree. We create trees for all the above mentioned methods for both the simulated and experimental datasets. We also compare with `Terminus` groups (`Term`) and transcripts(`Txp`) whenever possible. The total number of transcripts covered by different methods across the different datasets are provided in Table S2. `Mean` tree covers the most transcripts followed closely by `Cons` tree while `ConsFiltES` tree and `Term` cover the least number of transcripts. In order to facilitate the comparison between the methods, we ensure that all trees cover the same transcripts. A transcript set is created by taking the union of the transcripts covered by the trees in the set {`Mean`, `Cons`, `ConsFilt`, `ConsFiltES`, `AC}`, and groups in `Term`, with `Txp` referring to that transcript set. To the children of the root node of the above trees, we append any transcripts that were present in `Txp` but missing in the transcripts covered by the individual trees. Similarly, nodes in `Term` consist of all nodes output by `Terminus` along with any transcripts that are present in `Txp` but not covered by `Terminus` groups.

## 4. Results

### 4.1 TreeTerminus nodes have the lowest mean inferential variance

The violin plot in Figure 2 compares the distribution of log_2_ MIRV (mean inferential relative variance) for the inner nodes across trees stratified by their height for the `BrSimNorm` dataset. For comparison, log_2_ MIRV for genes and transcripts has been plotted along with the trees at each height. The transcripts have the highest range of variation in the mean inferential relative variance (MIRV) and as expected have the largest values. Genes have the lowest MIRV with the majority of genes having the lowest possible value of 0.01 that can be obtained using the default thresholds(Equation 2). Among the trees, the nodes of `Cons` and `Mean` trees have the lowest MIRV across different heights followed by `ConsFilt` and Anti-Correlation trees (`AC`). The nodes belonging to `ConsFiltES` tree have the highest MIRV. As expected, going up the tree, MIRV decreases across the methods getting closer to gene MIRV levels. These trends are summarized by the median values of MIRV in Table S3. The above pattern is also repeated in the `BrSimLow` dataset(Table S4, Figure S1). A similar trend is observed for the `MouseMuscle` dataset (Figure S2, Table S5). The only change is observed on the nodes at height greater than or equal to 5 for the `ConsFiltES` tree which shows the most downward shift in MIRV distribution. It is important to note that that across all the datasets, `ConsFiltES` has the lowest the number of nodes. For the `ChimpBrain` dataset, a very similar distribution is observed for the trees across all the methods and a modest downward shift in the upper tail is observed as the height increases. The MIRV though is very low for the height 2 inner nodes to begin with(Table S6, Figure S3). However, a considerable downward shift in the distribution of MIRV is still observed for the inner nodes compared to the transcripts.

**Fig. 2.**
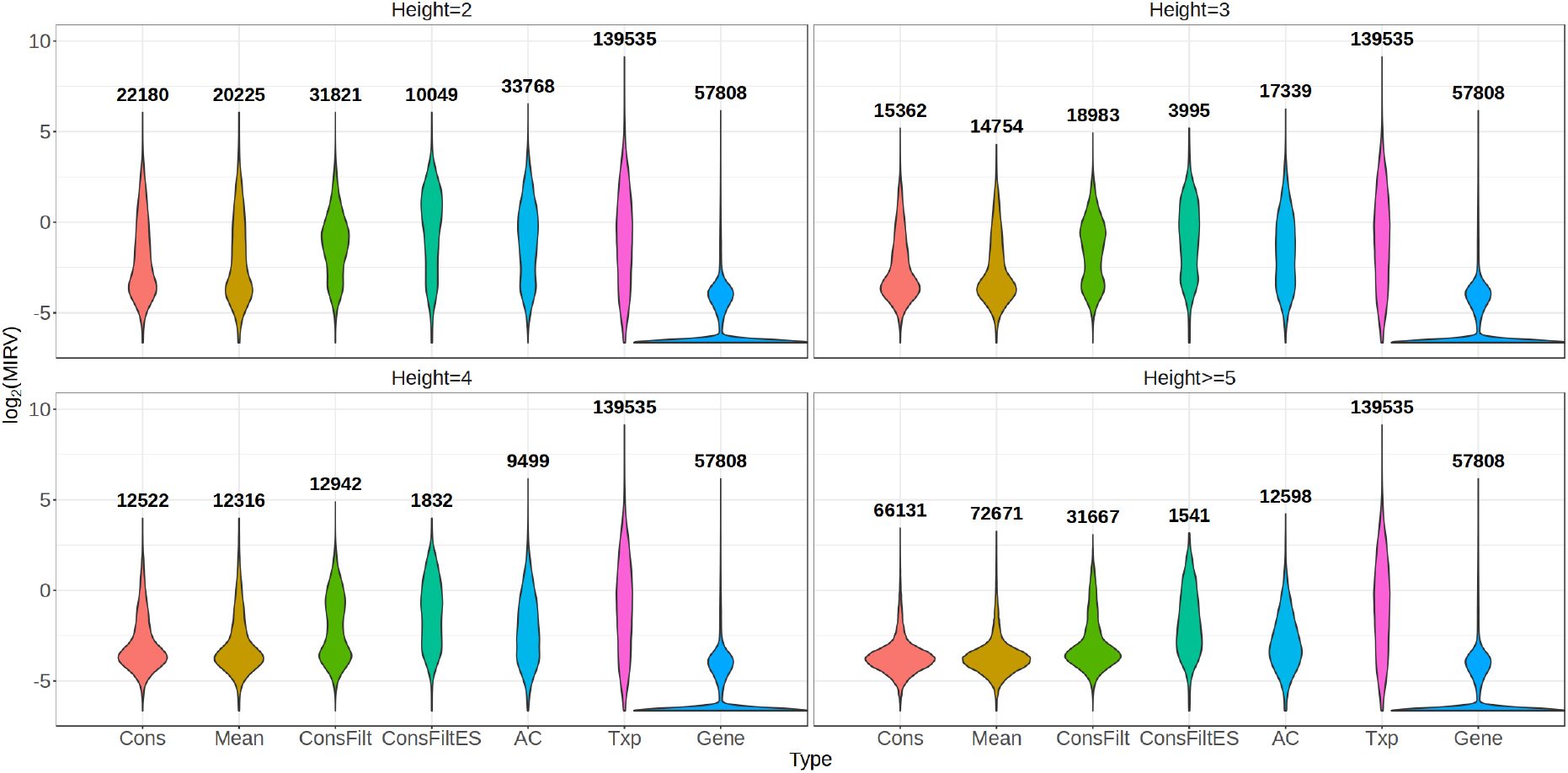
Distribution of log_2_ MIRV(mean inferential relative variance) across the inner nodes stratified by their height for different trees for the `BrSimNorm` Dataset, with the total number of inner nodes belonging to a method at a given height written on top of the violin plot. Also plotted for comparison at each height is the distribution of lg of MIRV for the transcripts and genes.

### 4.2 TreeTerminus nodes map to relatively fewer genes and gene families

We next look at the distribution of number of uniquely mapped genes at the inner nodes located at different heights across the trees. The mapped genes associated with a node are found by mapping the descendant leaves (transcripts) for that node to the genes. Figure 3, S4, S5 and S6 plots this distribution for the different datasets. The proportion of nodes that map to more than one gene is considerably higher for the `AC` tree compared to others, with the number of nodes at height two that map to two genes, tens to hundreds of times larger for the anti-correlation (`AC`) trees. The increase in the relative number of multi-gene mapping nodes is also the highest for the `AC` tree and it has the highest magnitude of the number of unique genes to which an inner node can map. With the exception of `AC` tree, the proportion of nodes that map to only one unique gene dominate nodes mapping to multiple genes across different heights in the other trees. The `AC` tree has the highest number of inner nodes that map to more than one hundred genes across the datasets (Table S7). No such nodes exist in the other trees obtained across the datasets, barring the `MouseMuscle` dataset, for which there exists a reasonable number of such inner nodes, with some even mapping to more than a thousand genes. When we looked at the genes mapping to these nodes, we found that a large proportion of such genes were either predicted or pseudogenes, which was not necessarily the case for most of such nodes in the `AC` tree.

**Fig. 3.**
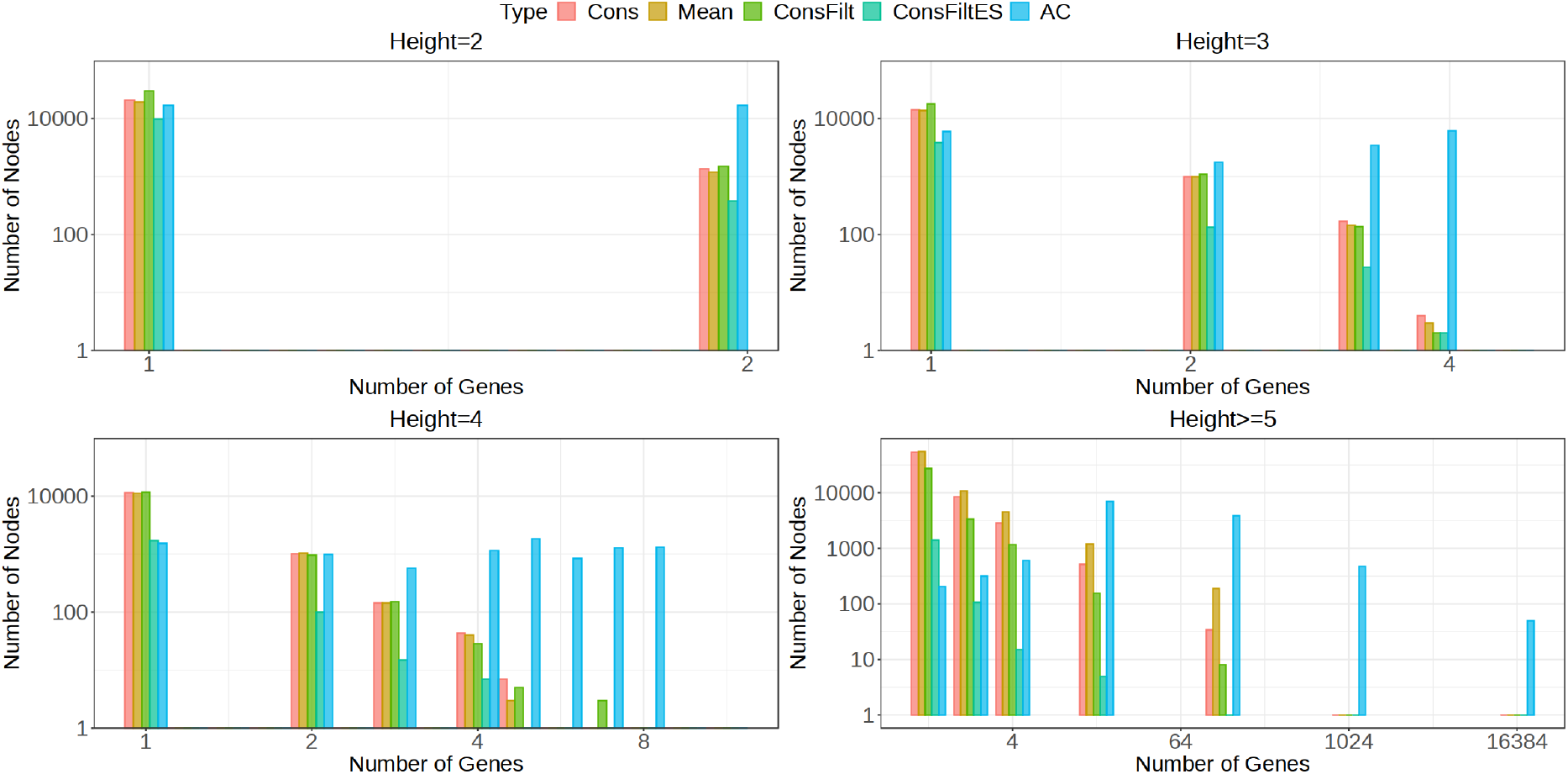
Comparison of different methods with respect to the number of genes to which an inner node in the tree maps for the `BrSimNorm` dataset, stratified by their height. The x-axis represents the number of unique genes that transcripts belonging to the inner nodes map to and y-axis represents the frequency of such mappings at a given height for a tree. For all the inner nodes located at a height greater than or equal to 5, number of unique genes were binned using the set {1,2,4,16,128,1024,16384}, with the bin representing number of unique genes less than or equal to the bin but larger than the bin left to it.

We also explored the distribution of the number of gene families to which inner nodes belonging to different trees map in Figure 4. The procedure for finding the gene families that map to a given inner node is described in S1.6. The number of gene families have been binned into bin numbers {1,2,4,10,100,500,1000,12000} so that each bin represents the number of unique gene families less than or equal to the bin but larger than the bin left to it. For the `MouseMuscle` dataset, the number of nodes mapping to more than one family is the largest for the `AC` tree and the difference in the number of nodes increases between `AC` and the remaining trees as the magnitude of number of gene families they map to is increased on the x-axis. While the `AC` tree has more than 5000 nodes that map to (10,100] gene families, this number is only 124 for the Mean tree and much less for the other trees. There are more than 500 nodes that map to more than 100 gene families for the `AC` tree while hardly any such node exists for the other trees. A similar trend is observed for the `ChimpBrain` dataset, while for the `AC` tree there are more than 2500 nodes that map to more than 10 gene families, such nodes don’t exist for the other trees. The nodes belonging to AC tree thus map to many more gene families. While the nodes belonging to the other trees map to a fewer gene families, we have not explored them in the current work.

**Fig. 4.**
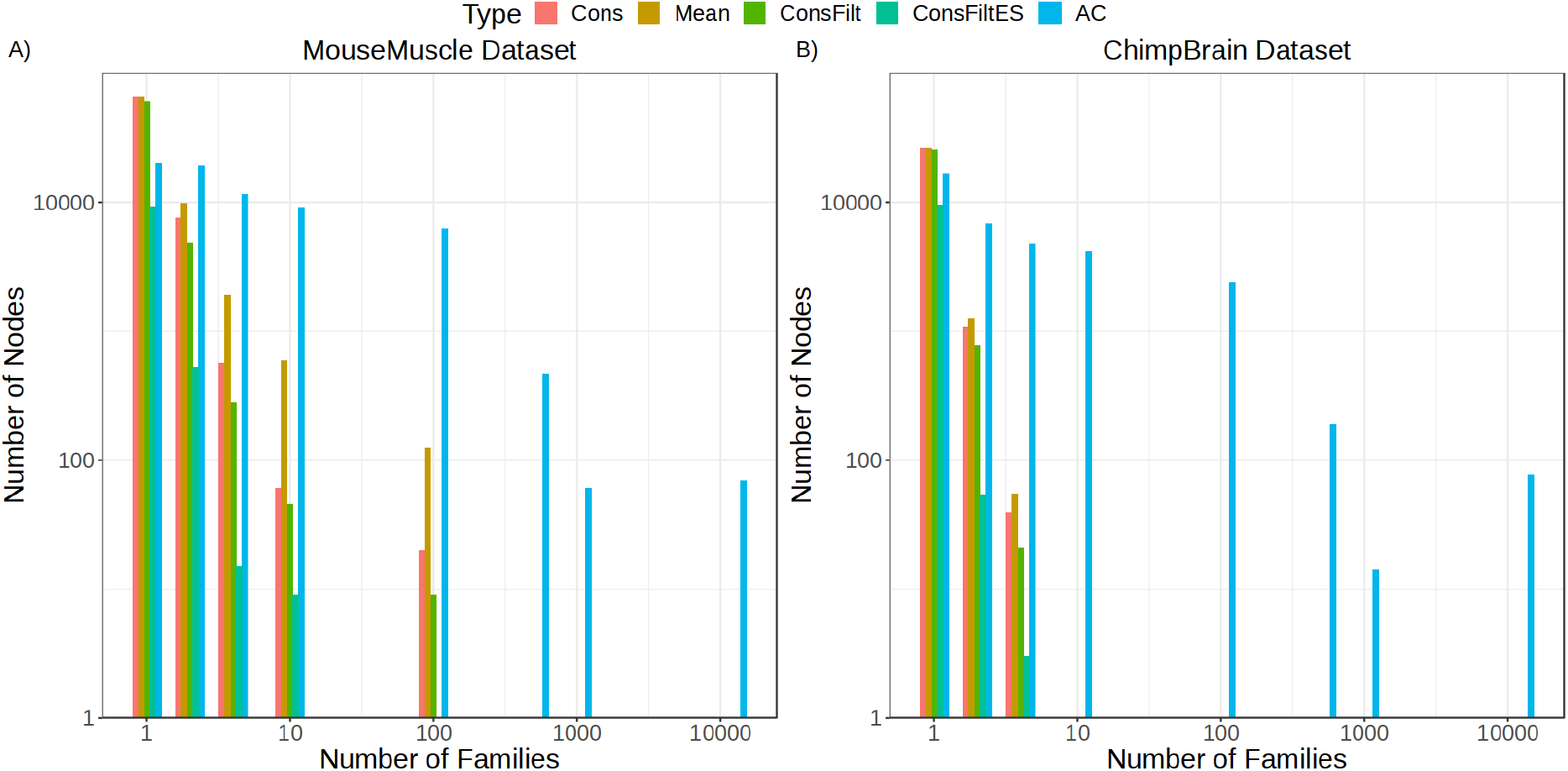
Comparison of different tree methods with respect to the number of gene families to which an inner nodes maps. Number of gene families that map to an inner node have been binned into the set {1,2,4,10,100,500,1000,12000}, with the bin representing the number of unique gene families less than or equal to the bin but larger than the bin left to it. The x-axis represents the bin and y-axis represents the total frequency of inner nodes mapping to the gene families in that bin for a given tree. This has been plotted for A) `MouseMuscle` Dataset and B) `ChimpBrain` Dataset.

### 4.3 TreeTerminus produces better optimal cuts for different objective functions

Figure 5, S7 plots the optimal values and the cut sizes obtained by solving for the objective function that minimizes the metric `irv height desc` (Section 2.4) on the different trees for each dataset. Also plotted for comparison are these values when the cut consists of only the transcripts (`Txp`). The cuts for different trees have been obtained by using *γ* values from the set {0.05,0.1,0.5,1,5}. The highest(worst) optimal value is seen for the cut consisting of transcripts, whereas the cuts obtained for `Cons`, `ConsFilt`, `Mean` trees consistently have the lowest(best) optimal values. The relative differences in the optimal value between them is smaller compared to differences in optimal values for the cuts obtained from the other trees. As *γ* increases, the size of the cut increases as expected along with the increase in the optimal value. Further, the differences in the optimal values and cut sizes between the methods also decreases, with these values individually becoming comparable at *γ* =5 across the trees. We also looked at the distribution of the values of the metric in the cuts for the different trees and the transcripts in Figure S8. A large proportion of nodes for the `Cons`, `Mean`, `ConsFilt` trees show a very low value for the metric compared to the other methods, showing that the net value of the objective function is not dominated by just a few outlier nodes.

**Fig. 5.**
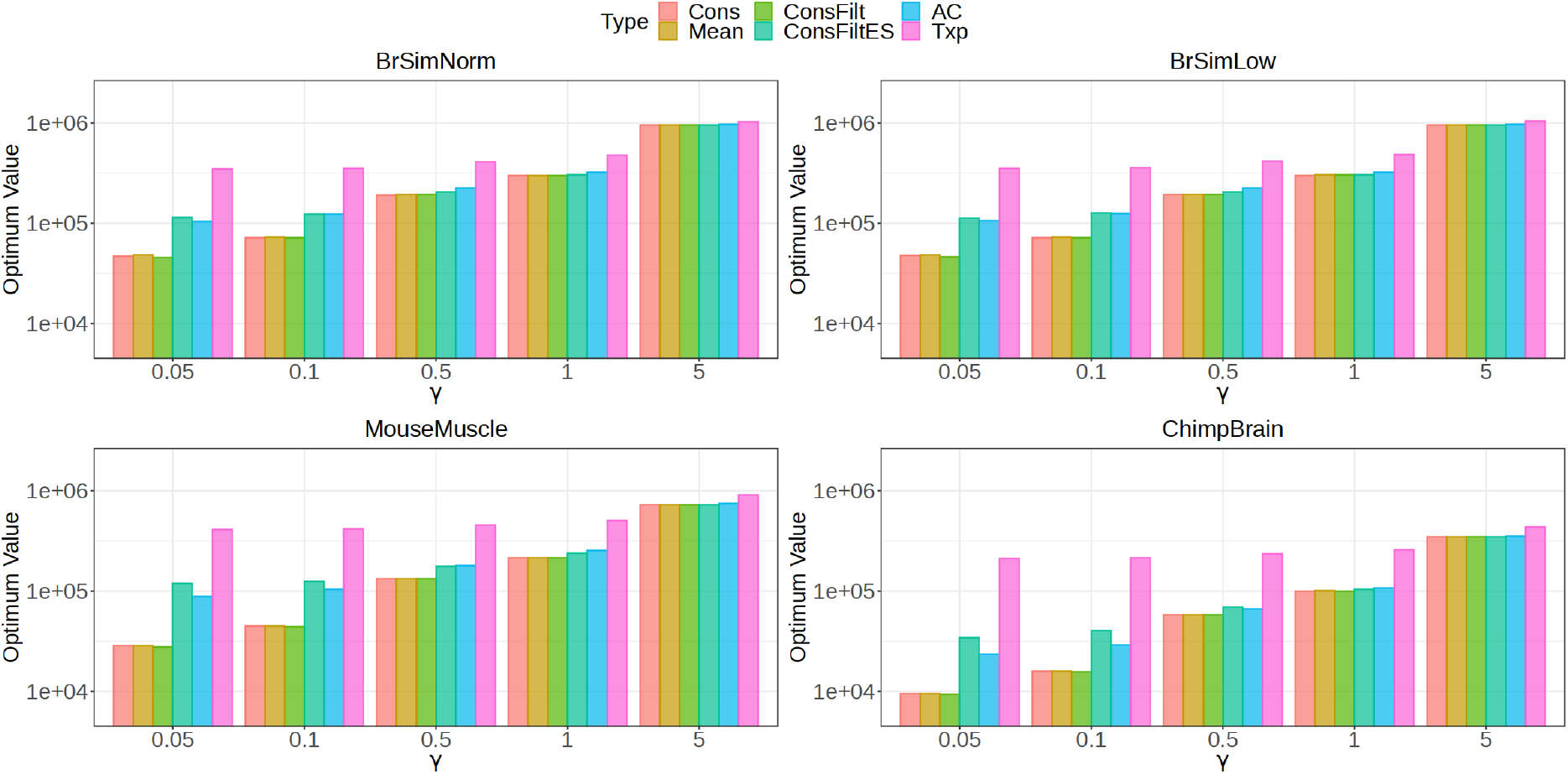
Barplot representing the values for the objective function on the cut obtained by minimizing the sum of metric -sum of the metric (`irv_height_desc`) on the different datasets. For each dataset, optimal values of the objective function are compared across the methods for a range of *γ* values. Also plotted for comparison is the optimal value obtained by using transcripts(`Txp`) as the cut.

We next compared the optimal values and the corresponding cut sizes, obtained by solving for the objective function that maximizes the sum of the metric `wlfc_desc` (see Section 2.4) for the different trees on each dataset in Figure S9. Also compared are these values that are obtained when the cut consists of - Terminus (`Term`) groups and the transcripts(`Txp`). The cuts obtained from `Mean` tree have the highest (best) optimal value followed closely by `Cons` tree. These are then followed by the cuts obtained for the `ConsFilt` and `AC` tree. The cuts obtained for `ConsFiltES` tree, `Term` and `Txp` have the lowest (worst) optimal values. An opposite trend is observed for the cut sizes, with `Txp` having the largest size followed by cuts from `Term`, and `ConsFiltES` tree. There was also a rightward shift in the distribution of the metric for the nodes obtained from the `Cons`,`Mean` trees, indicating that the increased optimal value for these trees was not due to the presence of just a few outlier nodes (Figure S10).

## 5 Discussion

`TreeTerminus` organizes the transcripts in a tree structure, with the leaves representing the transcripts. The tree accounts for experiment-wide inferential uncertainty and provides the flexibility to analyze data at nodes that are at different levels of resolution, that are best supported by data for the downstream analysis of interest. The cut given by the dynamic programming (DP) implementation on the objective functions used in this paper provides one possible way to get a set of nodes that can be used for downstream tasks. The DP is generalized and the base metric inside the objective function can be easily replaced by a user-defined function. Furthermore, the end user can also define a completely different approach to find a cut from the tree.

We have provided two different approaches for getting the transcript trees as output from an RNA-Seq experiment, `Mean` and `Cons` trees. They both cover a similar number of transcripts and have comparable performance across the different datasets for the various analysis that have been explored in this paper. They have a superior performance compared to the consensus trees that were obtained when the constraints were kept on the various evaluation metrics and cover the largest number of transcripts. The sub-par performance for the other methods, especially `ConsFiltES` and `Term` perhaps indicate that these methods under-aggregate. While no other method exists that arrange the transcripts in a tree-like structure, for comparison we also provide the `AC` tree as a baseline, that is constructed using UPGMA - a well known method for tree construction in phylogenetics. However, construction of the `AC` tree is both highly memory and time intensive, with the tree construction process not taking into account the equivalence class information. As a result, we observe that the nodes belonging to `AC` tree map to a large number of distinct genes and gene families, not offering much biological interpretation. The nodes belonging to trees constructed from `TreeTerminus` map to a smaller number of distinct gene and gene families, even though annotation of the underlying organism was not a part of the input.

There do exist some areas where the underlying tree constructions methods can be improved further. Taking the mean of difference in inferential relative variance on all samples for constructing `Mean` tree might prevent transcripts from aggregation that had high uncertainty but were expressed only in the samples belonging to a population that was not in the majority in the experiment. This can be replaced by weighted reduction in inferential relative variance, where the mean is computed over the samples in which the transcripts are expressed. A limitation with the `Cons` tree is the consensus tree algorithm that requires all the underlying input trees to span the same leaf set, causing us to modify the individual sample group trees that were provided as an input. An alternative to consensus algorithms can be supertree methods such as STELAR [18] and FastRFS [45], which don’t require that all the input trees should cover the same leaf set. However, STELAR is very slow for trees that span large leaf sets and FastRFS provides an unrooted tree as an output, which makes using them as a direct replacement not trivial. Modifications to these approaches can be explored in the future. Differential expression or differential abundance analysis is an important downstream application of RNA-Seq and other sequencing-based datasets. Various differential testing methods that leverage a tree structure [16, 25, 4, 29] have been proposed, reporting an increased power. In microbiome analysis, investigators have tried to associate nodes, located at different levels on the tree built through phylogenetic analysis, with a response variable of interest[5], with some using replicability as a metric to determine optimal levels of aggregation[9]. We believe that the tree(unified) provided by `TreeTerminus` can be used with these methods for an improved differential analysis in RNA-Seq. Since none of the methods have been designed taking RNA-Seq analysis into account, there lies an opportunity to develop tree differential testing methods for RNA-Seq data that output nodes that are at the finest resolution and at the same time have a robust signal. Further, `TreeTerminus` in the future can also be extended to tagged-end scRNA-seq protocols where the trees will be constructed on genes rather than transcripts, owing to read mapping ambiguity at the gene level itself since the exons of one gene can overlap with exons or introns of other genes [38], as these are 3’-biased, often single end reads.

