## Supplementary Material for "TreeTerminus - Creating transcript trees using inferential replicate counts"

### S1 Methods

#### S1.1 Terminus

**Consensus** - The consensus procedure outputs a set of transcript groups across samples. The procedure involves creating a connected undirected graph on the transcripts, with an edge between any pair of transcripts denoting that they co-occur in a group in any sample and the edge weight denotes the count of the total number of samples in which the pair co-occurs in a group. To create the consensus groups, it finds the connected components on the updated graph obtained after removing the edges that occur in atleast a certain proportion of samples. This proportion is a user-defined parameter and by default has been set to 0.5.

#### S1.2 Majority rule extended consensus tree

We borrow the notation from [19], to define a consensus tree. Given a tree $T$, let $V(T)$ denote the set of all nodes and $\Lambda(T)$ denotes the set of transcripts covered by the tree $T$. For any node $u \in V(T)$, $T[u]$ is the subtree rooted at $u$, $\Lambda(T[u])$ is the cluster associated with it and $C(T)$ represents the set of all clusters associated with tree $T$. Let $S$ denote the collection of trees $(T_1, T_2, ..T_M)$, with $\Lambda(T_i) = L, \forall i \in \{1, ..., M\}$, aka all trees have the same leaf set. Let $X$ be the set of all clusters that occur in $S$ sorted by the decreasing order of the frequencies with which they occur in the trees. Construct a set $Y$ of clusters as:
Initialize $Y = \emptyset$, then traverse $X$ and for each cluster $C$ encountered in this order, check if $C$ and $C'$ are pairwise compatible for all $C' \in Y$, if yes then $Y = Y \cup \{C\}$. A greedy or majority extended consensus tree of $S$ is a tree $T$ such that $\Lambda(T) = L$ and $C(T) = Y$.

#### S1.3 Construction of anti-correlation tree

All transcripts that had 0 counts across the samples and 0 counts across the inferential replicates for any sample were removed. For each sample $k$, Pearson correlation $r_{ij}^k$ was computed between each pair of transcripts across the inferential replicates. $r_{ij}^k$ is multiplied by $-1$ to give ${r_{ij}^k}'$ so that any pair of transcripts that had the highest negative correlation now have the largest positive correlation. To convert (anti)correlation into distance, ${r_{ij}^k}'$ is transformed as $d_{ij}^k = (1 - {r_{ij}^k}')/2$ or $d_{ij}^k = \frac{1+r_{ij}^k}{2}$. Unweighted Pair Group Method with Arithmetic Mean (UPGMA) [37] was used to create the anti-correlation tree with the implementation derived from the `R` package `phangorn` [34]. The mean for $d_{ij}^k$ across samples is computed and fed as input to UPGMA in order to get the final tree. The `AC` tree could also have been created for every sample and then provided as input to the consensus tree algorithm to get the final tree, however consensus tree algorithms do not scale well on the number of leaves. Since for most cases the number of leaves(transcripts) on trees would be in the order $10^4 - 10^5$, consensus tree algorithms would have taken a lot of time to converge.

#### S1.4 Datasets

To demonstrate the benefits of TreeTerminus, we ran it on both simulated and experimental datasets spanning different organisms.

**Simulated human datasets** Polyester[14] was used to generate simulated RNA-seq data. TPM estimates were extracted from GTEx V8 frontal cortex dataset with the distribution of mean and dispersion values derived from GEUVADIS samples [22]. The process of read generation has been described in detail in [26]. We generated 12 samples with 6 samples in each condition. All transcripts were differentially expressed for 10% of the genes with all having the same fold change (DGE) and 10% genes had a single transcript differentially expressed (DTE). We created two variations of this simulation by varying the range of fold change - in the first variation we keep the same fold change as in [26] and in the second variation the maximum range of fold change is lowered from 6 to 3 . The first variation is referred to as BrSimNorm and second is referred as BrSimLow.

**Mouse muscle dataset** This dataset is taken from the skeletal muscle study GSE100505[42]. In this paper, we have used 12 samples with 6 samples belonging to Atria and 6 to Tibialis Anterior, with the accession numbers provided in Table S1. All the samples belong to organism mus musculus. This dataset is referred as MouseMuscle.

**Chimpanzee brain dataset** The final dataset that has been analyzed in our study is the RNA-Seq data from [39], SynapseID - syn7067053 collected from 5 Chimpanzees (Pan Troglodyte). We refer to this dataset as ChimpBrain. For each specimen, samples from 16 different tissues representing hippocampus, amygdala, cerebellar cortex, mediodorsal nucelus of thalamus, striatum and 11 areas of neocortex were sequenced. The samples belonging to the medial dorsal nucleus were removed before running TreeTerminus. The lfc change was computed by taking 5 cerebellum samples as the first group and remaining 68 samples belonging to other tissues as the second group. Batch effects were observed w.r.t specimen label and corrected using sva[23].

#### S1.5 Quantification pipeline

For the Chimpanzee dataset, only bam files were available as the raw data which were converted into fastq using bamToFastq in bedtools [31]. The quality control analysis for Chimpanzee and mice dataset was done using fastqc[2] and multiqc[10]. For creating the salmon indexes, gencode versions v26, vM25 and Pan_tro 3.0 were used for human, mice and chimpanzee datasets. Salmon was used for quantification and generating 100 Gibbs replicates for each sample with a thinning factor of 100. All the pipelines used for analysis in this paper were created using Snakemake [21].

#### S1.6 Mapping to gene families

The gene family labels for a gene were extracted using R package biomaRt [36] using Ensembl version 101. We want to extract the total number of unique gene families to which an inner node maps. A gene can map to more than one gene family and when an inner node maps to multiple genes; with atleast one gene mapping to more than one family, finding the unique number of gene families for that node is not trivial. Simply, taking a union of gene families across genes for an inner node is undesirable since this might lead to a situation where number of reported gene families are larger than the number of mapped genes for some nodes. Between different trees, the interpretation of the distribution of the number of gene families to which an inner node maps will also be biased towards the tree that has more nodes containing genes mapping to multiple gene families. Thus, in order to find the number of unique gene families associated with a node, we formulate this as the minimial hitting set problem [20]. We want to find the minimum number of gene families for a node in the tree, whose intersection with the gene families associated with every gene for that node leads to a non-empty set. We use the implementation provided by the Python library PySAT [17] to solve the minimal hitting set problem.

### S2 Tables

**Table S1.** Accession IDs for the tissue samples for the `MouseMuscle` dataset that have been used for analysis in this paper

| **Accession ID** | **TissueName** |
|---|---|
| SRR5758624 | Atria |
| SRR5758625 | Atria |
| SRR5758626 | Atria |
| SRR5758627 | Atria |
| SRR5758628 | Atria |
| SRR5758629 | Atria |
| SRR5758707 | TA |
| SRR5758706 | TA |
| SRR5758705 | TA |
| SRR5758704 | TA |
| SRR5758703 | TA |
| SRR5758702 | TA |

**Table S2.** Total number of transcripts covered by different methods across the datasets.

| **Method** | **BrSimNorm** | **BrSimLow** | **MouseMuscle** | **ChimpBrain** |
|---|---|---|---|---|
| Mean | 135138 | 135208 | 98091 | 39856 |
| Cons | 134377 | 134371 | 97360 | 39491 |
| ConsFilt | 121784 | 121657 | 89429 | 37858 |
| AC | 73205 | 73493 | 68062 | 35114 |
| ConsFiltES | 27156 | 26891 | 17939 | 15773 |
| Term | 13612 | 13730 | 7792 | 5428 |

**Table S3.** Median of mean inferential variance (MIRV) of the inner nodes for different trees stratified by their height for the `BrSimNorm` dataset. All nodes with height larger than 5 have been labelled as 5.

| **Tree** | **2** | **3** | **4** | **5** |
|---|---|---|---|---|
| Mean | 0.17 | 0.11 | 0.09 | 0.07 |
| Cons | 0.19 | 0.12 | 0.10 | 0.07 |
| AC | 0.52 | 0.32 | 0.21 | 0.12 |
| ConsFilt | 0.44 | 0.35 | 0.19 | 0.10 |
| ConsFiltES | 0.89 | 0.56 | 0.42 | 0.26 |

**Table S4.** Median of mean inferential variance (MIRV) of the inner nodes for different trees stratified by their height for the `BrSimLow` dataset. All nodes with height larger than 5 have been labelled as 5.

| **Tree** | **2** | **3** | **4** | **5** |
|---|---|---|---|---|
| Mean | 0.17 | 0.11 | 0.09 | 0.07 |
| Cons | 0.19 | 0.12 | 0.10 | 0.07 |
| AC | 0.52 | 0.33 | 0.21 | 0.13 |
| ConsFilt | 0.44 | 0.35 | 0.19 | 0.10 |
| ConsFiltES | 0.90 | 0.57 | 0.42 | 0.30 |

**Table S5.** Median of mean inferential variance (MIRV) of the inner nodes for different trees stratified by their height for the `MouseMuscle` dataset. All nodes with height larger than 5 have been labelled as 5.

| **Tree** | **2** | **3** | **4** | **5** |
|---|---|---|---|---|
| Mean | 0.12 | 0.09 | 0.08 | 0.07 |
| Cons | 0.12 | 0.09 | 0.08 | 0.07 |
| AC | 0.47 | 0.35 | 0.24 | 0.19 |
| ConsFilt | 0.24 | 0.14 | 0.10 | 0.08 |
| ConsFiltES | 0.68 | 0.54 | 0.29 | 0.02 |

**Table S6.** Median of mean inferential variance (MIRV) of the inner nodes for different trees stratified by their height for the `ChimpBrain` dataset. All nodes with height larger than 5 have been labelled as 5

| **Tree** | **2** | **3** | **4** | **5** |
|---|---|---|---|---|
| AC | 0.09 | 0.07 | 0.06 | 0.06 |
| Cons | 0.08 | 0.07 | 0.07 | 0.07 |
| ConsRed | 0.09 | 0.07 | 0.07 | 0.07 |
| ConsRedInd | 0.37 | 0.17 | 0.12 | 0.10 |
| Mean | 0.07 | 0.07 | 0.07 | 0.07 |

**Table S7.** Total number of inner nodes mapping to more than 100 genes for different datasets.

| **Tree** | **BrSimNorm** | **BrSimLow** | **MouseMuscle** | **ChimpBrain** |
|---|---|---|---|---|
| Mean | 0 | 0 | 113 | 0 |
| Cons | 0 | 0 | 44 | 0 |
| ConsInd | 0 | 0 | 37 | 0 |
| AC | 717 | 719 | 696 | 304 |
| ConsRedInd | 0 | 0 | 36 | 0 |

### S3 Figures

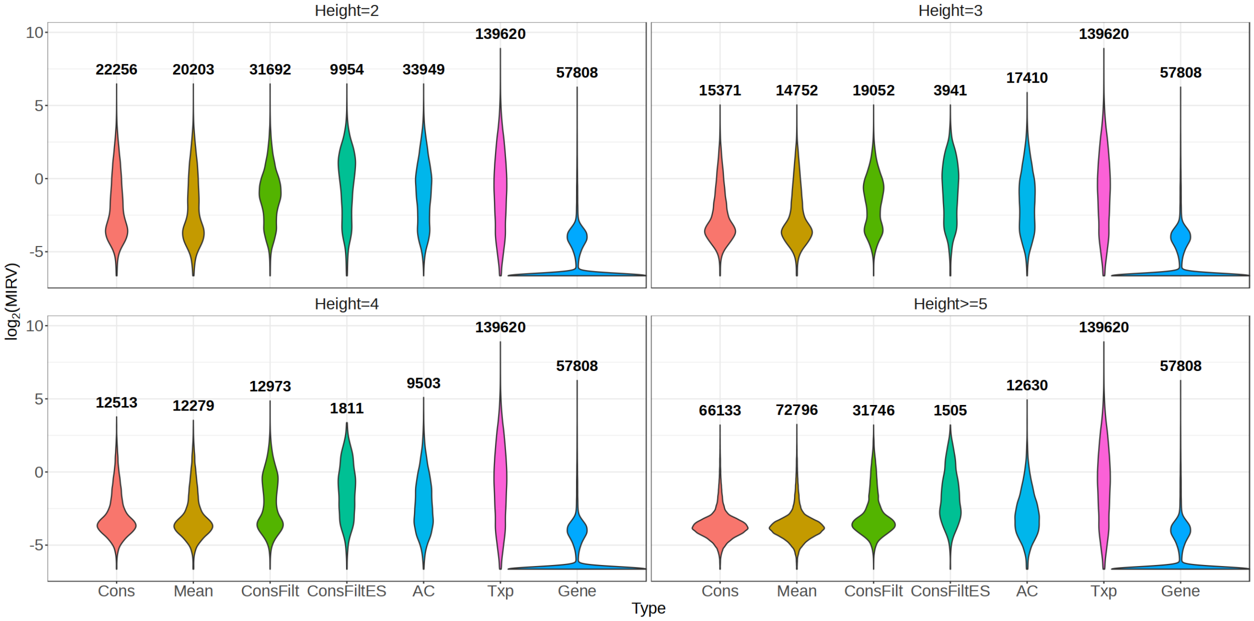


**Fig. S1.** Distribution of $\log_2$ MIRV (mean inferential variance) across samples for the inner nodes stratified by their height for different trees for the `BrSimLow` Dataset, with the total number of inner nodes belonging to a method at a given height written on top of the violin plot. Also plotted for comparison at each height is the distribution of MIRV for the transcripts and genes.

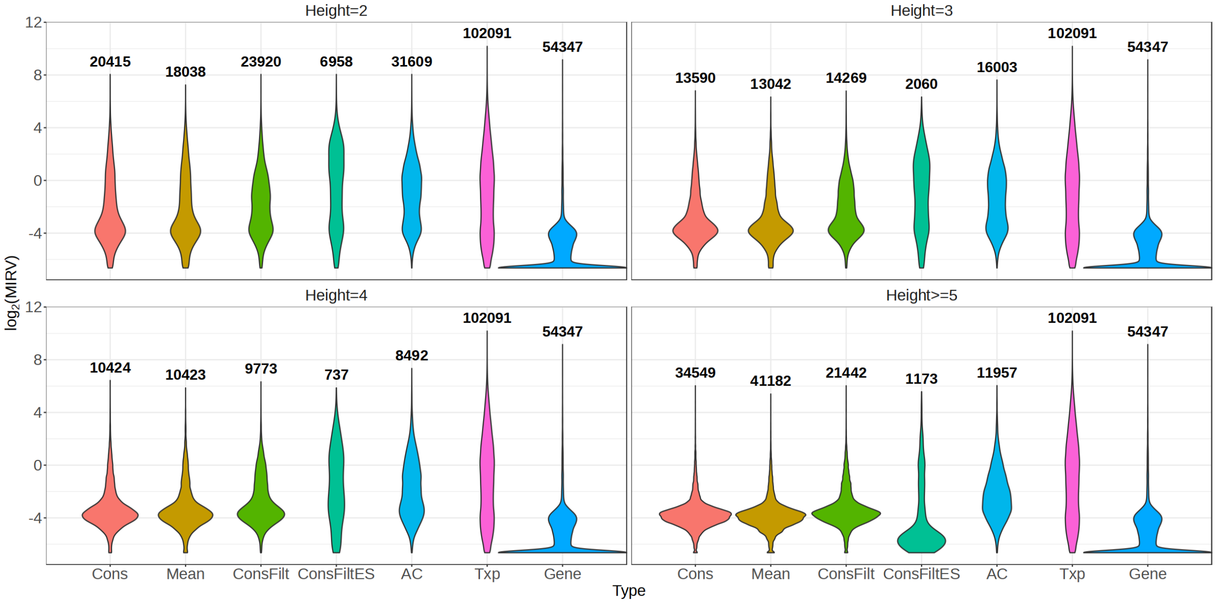


**Fig. S2.** Distribution of $\log_2$ MIRV (mean inferential variance) across the inner nodes stratified by their height for different trees for the `MouseMuscle` Dataset, with the total number of inner nodes belonging to a method at a given height written on top of the violin plot. Also plotted for comparison at each height is the distribution of lg of MIRV for the transcripts and genes.

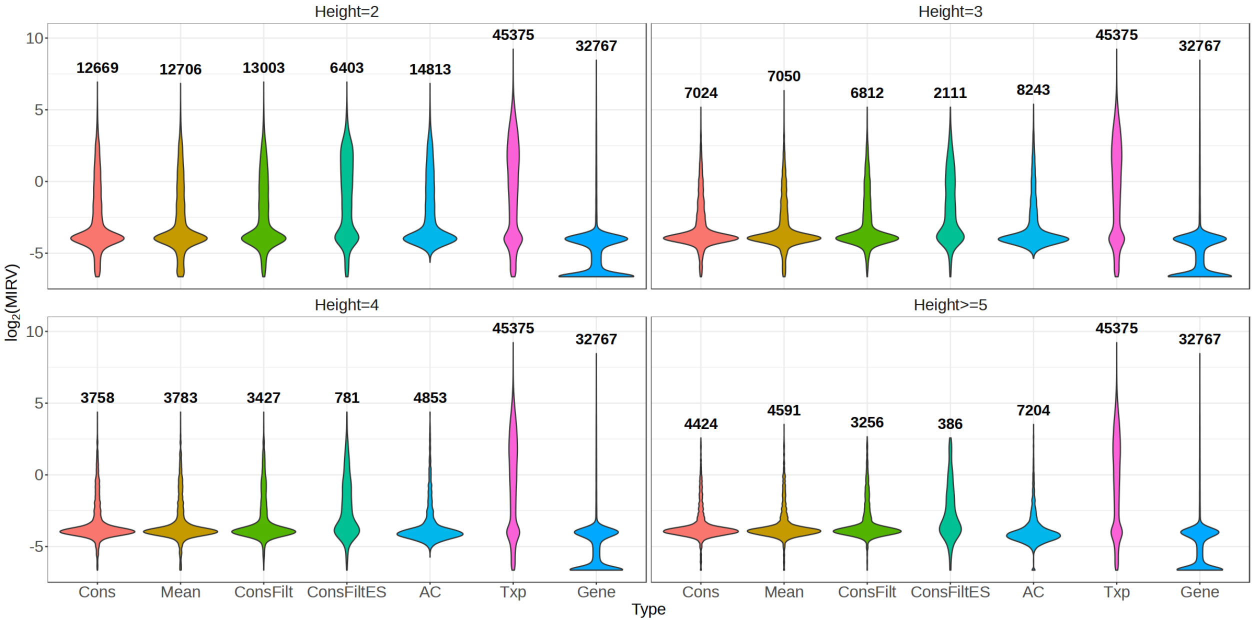


**Fig. S3.** Distribution of $\log_2$ MIRV (mean inferential variance) across samples for the inner nodes stratified by their height for different trees for the `ChimpBrain` Dataset, with the total number of inner nodes belonging to a method at a given height written on top of the violin plot. Also plotted for comparison at each height is the distribution of MIRV for the transcripts and genes.

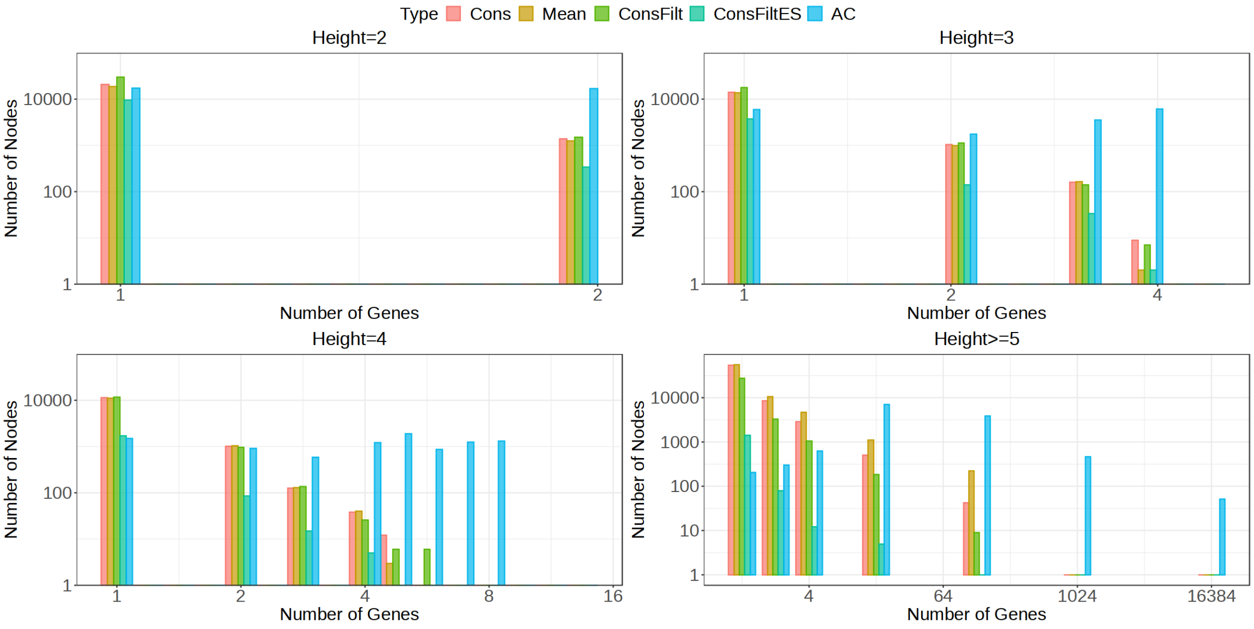


**Fig. S4.** Comparison of different tree methods with respect to the number of genes to which an inner node in the tree maps for the `BrSimLow` dataset stratified by their height. The x-axis represents the number of unique genes that transcripts belonging to the inner nodes map to and y-axis represents the frequency of such mappings at a given height for a tree. For all the inner nodes located at a height greater than or equal to 5, number of unique genes were binned using the set {1,2,4,16,128,1024,16384}, with the bin representing number of unique genes less than or equal to the bin but larger than the bin left to it.

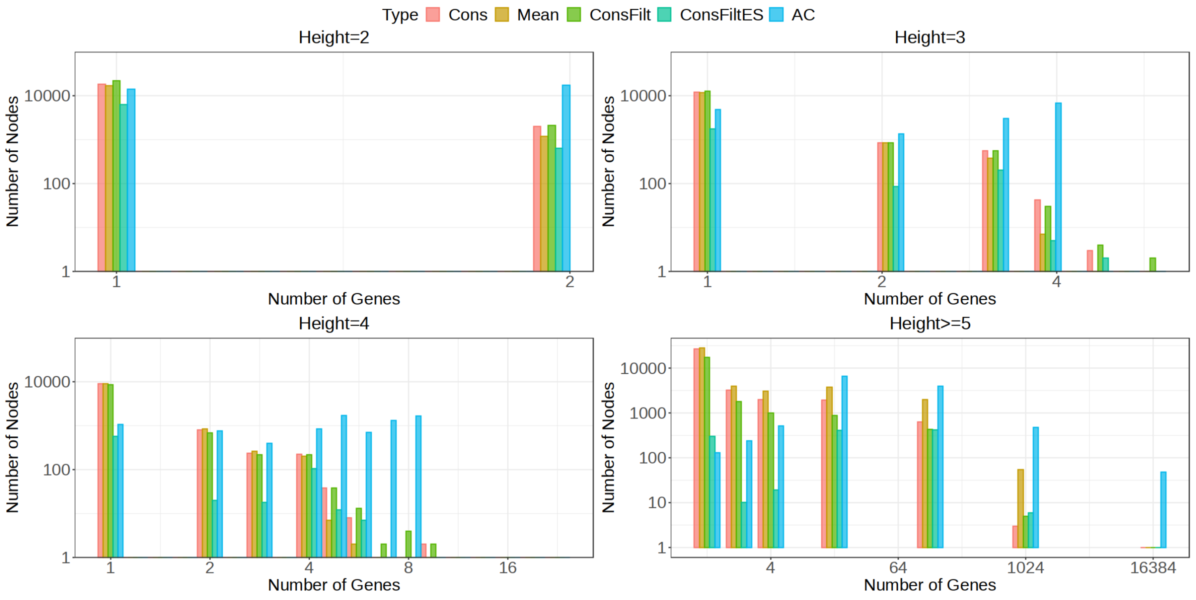


**Fig. S5.** Comparison of different tree methods with respect to the number of genes an to which an inner node in the tree maps for the `MouseMuscle` dataset stratified by their height. The x-axis represents the number of unique genes that transcripts belonging to the inner nodes map to and y-axis represents the frequency of such mappings at a given height for a tree. For all the inner nodes located at a height greater than or equal to 5, number of unique genes were binned using the set {1,2,4,16,128,1024,16384}, with the bin representing number of unique genes less than or equal to the bin but larger than the bin left to it.

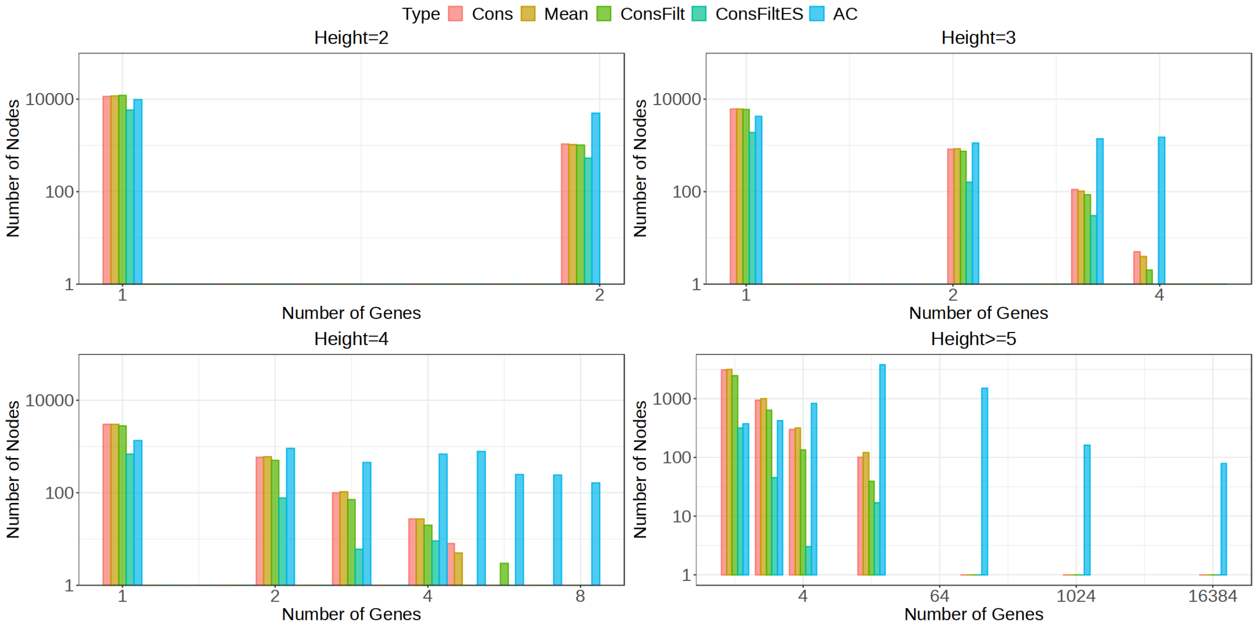


**Fig. S6.** Comparison of different tree methods with respect to the number of genes to which an inner node in the tree maps for the `ChimpBrain` dataset stratified by their height. The x-axis represents the number of unique genes that transcripts belonging to the inner nodes map to and y-axis represents the frequency of such mappings at a given height for a tree. For all the inner nodes located at a height greater than or equal to 5, number of unique genes were binned using the set {1,2,4,16,128,1024,16384}, with the bin representing number of unique genes less than or equal to the bin but larger than the bin left to it.

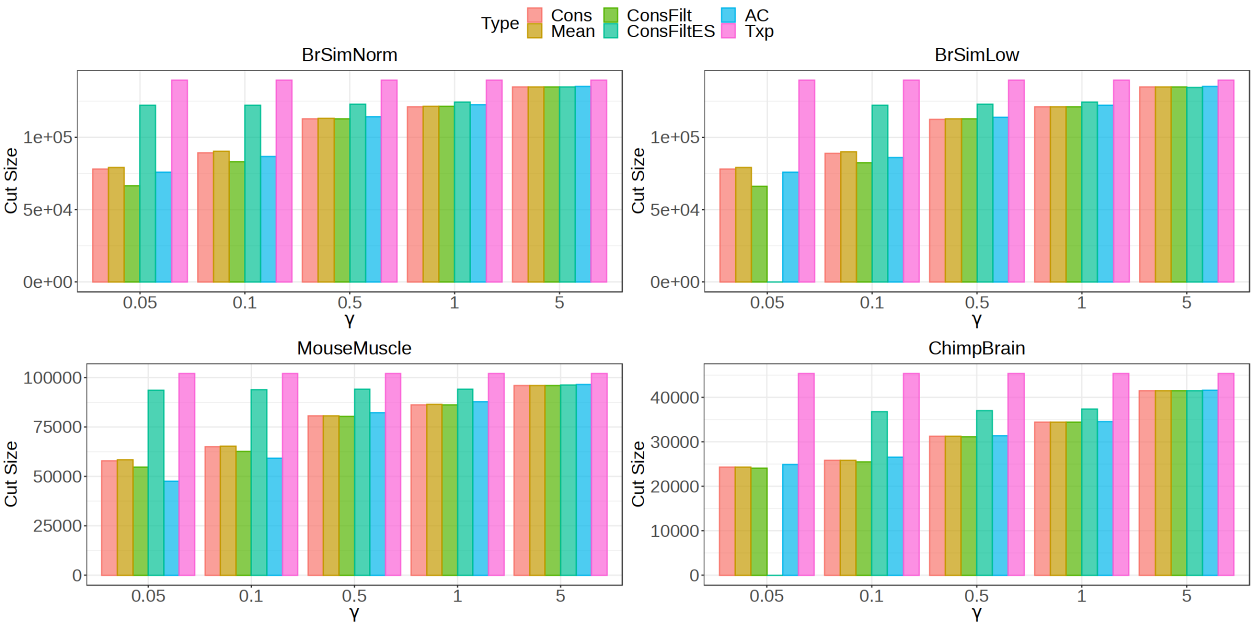


**Fig. S7.** Distribution of the size of cuts on the different datasets obtained after solving for the objective function that minimizes the sum of metric - weighted sum of mean inferential variance and height multiplied by the number of descendant leaves (`irv_height_desc`) for the nodes in a cut. For each method, the distribution is plotted for a range of $\gamma$ values. Also plotted for comparison are the total number of transcripts/leaves `Txp`.

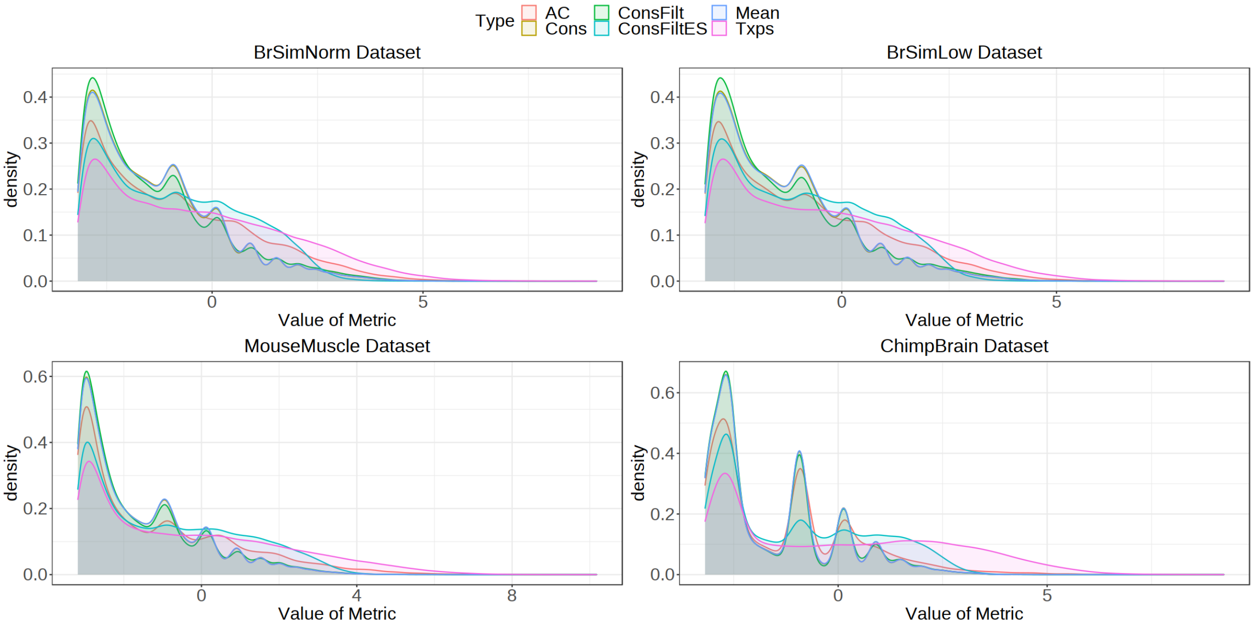


**Fig. S8.** Distribution of the lg of the metric - weighted sum of mean inferential variance and height multiplied by the number of descendant leaves (`irv_height_desc`) for the nodes in the cut obtained after minimizing for the objective function using `irv_height_desc` as the underlying metric across trees on the different datasets. The metric has been computed using $\gamma=0.1$. Also plotted for comparison is the distribution of `irv_height_desc` for transcripts.

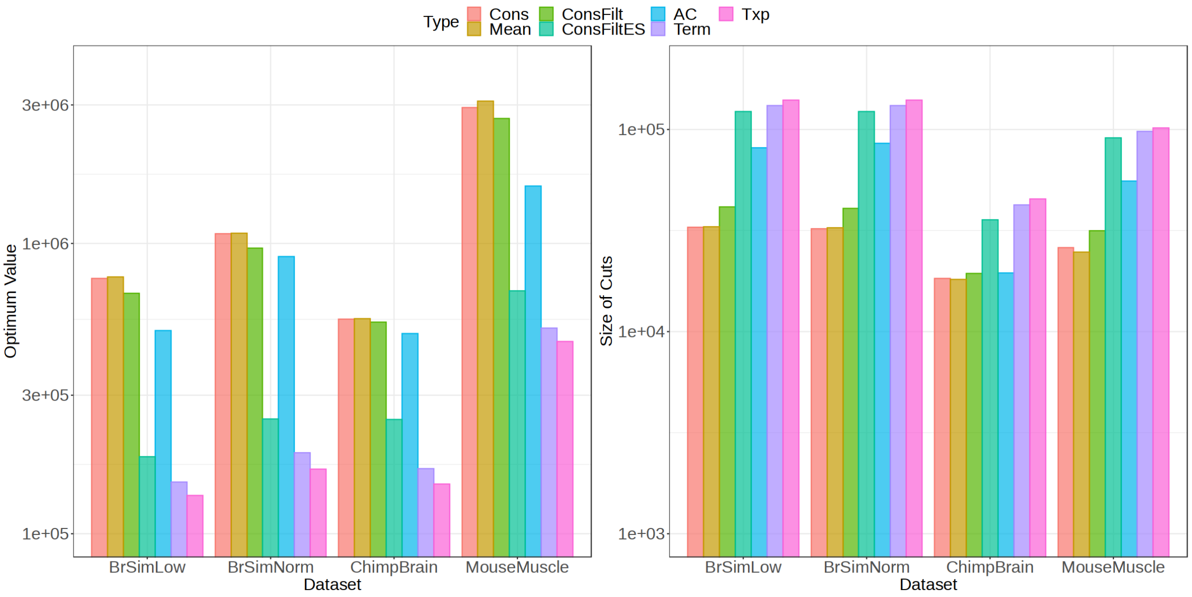


**Fig. S9.** Comparing the performance on the cuts across different trees obtained by solving the objective function that maximizes the sum of metric log fold change weighted by mean inferential relative variance (MIRV) multiplied by the number of descendant leaves (`wlfc_desc`) for the nodes in a cut on the different datasets. The performance is also compared when the transcripts and `Terminus` groups are taken as the cut. A) Value of the Objective Function. B) Size of the cut.

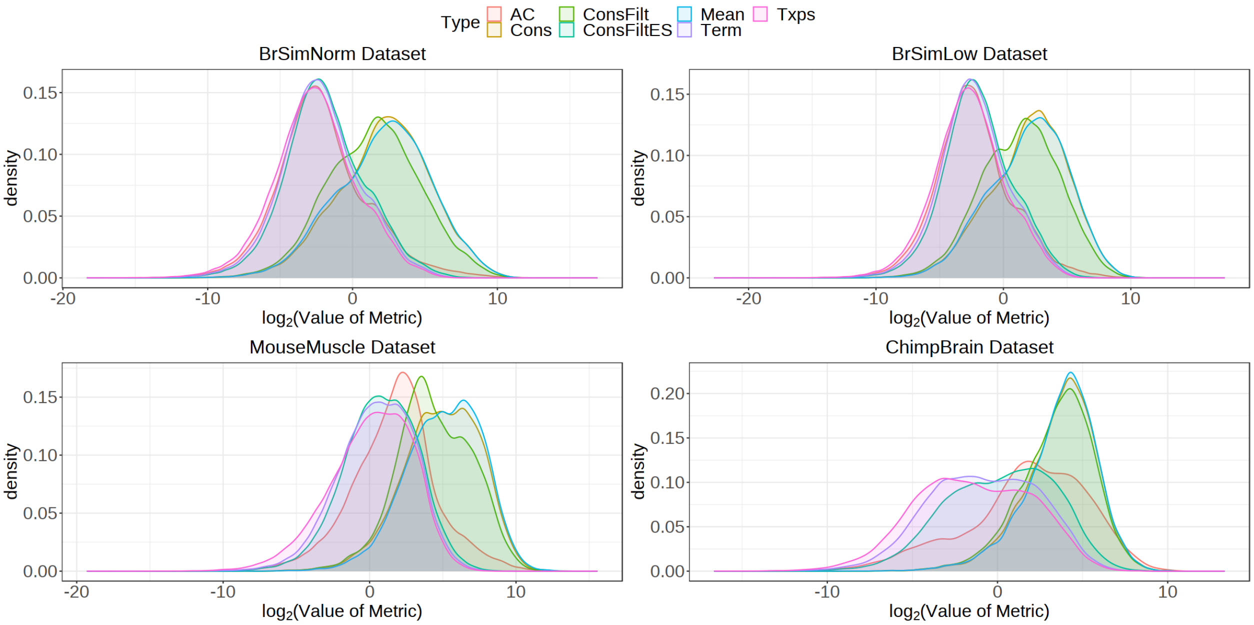


**Fig. S10.** Distribution of the lg of the metric - weighted log fold change multiplied by the number of descendant leaves (`wlfc_desc`) for the nodes in the cut obtained after maximizing for the objective function using `wlfc_desc` as the underlying metric across trees on the different datasets. Also plotted for comparison is the distribution of `wlfc_desc` for transcripts and `Terminus`.
